# Ontology-based workflow extraction from texts using word sense disambiguation

**DOI:** 10.1101/082784

**Authors:** Ahmed Halioui, Petko Valtchev, Abdoulaye Baniré Diallo

**Affiliations:** Département d’informatique, Université du Québec à Montréal (Québec), Canada.

**Keywords:** Workflow extraction, Ontologies, Word Sense Disambiguation, Similarity

## Abstract

This paper introduces a method for automatic workflow extraction from texts using Process-Oriented Case-Based Reasoning (POCBR). While the current workflow management systems implement mostly different complicated graphical tasks based on advanced distributed solutions (*e.g.* cloud computing and grid computation), workflow knowledge acquisition from texts using case-based reasoning represents more expressive and semantic cases representations. We propose in this context, an ontology-based workflow extraction framework to acquire processual knowledge from texts. Our methodology extends classic NLP techniques to extract and disambiguate tasks in texts. Using a graph-based representation of workflows and a domain ontology, our extraction process uses a context-based approach to recognize workflow components : data and control flows. We applied our framework in a technical domain in bioinformatics : i.e. phylogenetic analyses. An evaluation based on workflow semantic similarities on a gold standard proves that our approach provides promising results in the process extraction domain. Both data and implementation of our framework are available in : http://labo.bioinfo.uqam.ca/tgrowler.

## 1. Introduction

Current Workflow Management Systems provide a step by step guidance for users through sophisticated graphical workflow modelling [1]. Tracking these executions can be used to adapt workflows to new situations. This can be supported by reasoning methods like Process Oriented Case-Based Reasoning (POCBR) [2] which is a newly merged branch from the Case Based Reasoning (CBR) field. Because case engineering is a complicated and a costly process, automatic case-based acquisition from texts has been a major focus research in the last decade [3, 4, 5].

Textual CBR approaches vary widely in their scope according to their use of Natural Language Processing (NLP) methods and highly dependent on the intended applications. Even though [6] compared three different methods : key-based, grammar-based and machine learning, they considered only imperative sentence which cover more than 50% of the whole data and their interpretation modules are limited to the command tuple representation (A : type of the action, textual parameter to an action, the target on which to act). In [7], the authors developed a process extractor trained with a few manually created positive samples. A set of extraction rules are generated and then applied to automatically crawled documents (from e-how web sites, e.g. www.wikihow.com). However, their model have only successfully extracted ~ 42% of these documents. In [4], the authors propose to use a full fledged NLP solution to extract and enrich workflows from texts. Their frame-based Information Extraction (IE) approach to resolve anaphoras in texts uses *frame* patterns to trigger phrases with slots filled with content extracted from texts. A sequential pattern mining is also applied to mine anaphora-rules and then search for the most frequent term cooccurrences to resolve an anaphoric reference. A similar approach is proposed by [3] to resolve "universal references" using a domain-specific ontology. Given that ontology, they map the anaphoric reference to a concept. Their resolution is based on heuristics and very dependent to the domain application. In [8], the authors proposed a Plan Acquisition Architecture to build an augmented plan in the form of predicates from extracted raw content by reducing the article into a list of two kinds of information : input-tool (predicate that describes the tool properties) and action (action description with action properties). They also use data mining and statistical algorithms to identify particular subsequences of plans to specific contexts. Instead of manually composed plans from a set of atomic actions, the authors in [9] proposed to generate plans by transforming natural language task instructions from how-to sites into formal symbolic robot plans. Their evaluation results show that their planner is able transform about 80% of the instructions. This number can be increased with a better syntax parser and more mappings to find connections between WordNet synsets and ontological concepts. The most recent approach to extracting situation ontology from how-to instructions is found in [10] where an automatic method using syntactic patterns and CRF-based probabilistic model are used to classify actions and objects such as time and place. Although the pattern-based approach achieves a high level accuracy, its coverage is limited to the rules constructed semi-automatically from small fraction of sentences and possible patterns to reveal the associations between the verb and the subject/object. In [11], the authors proposed a novel attempt to extract procedural knowledge from Medline abstracts using the TAM model — a triple combination of Target, Method, and Action, and designed a text mining method with deep language processing analysis. This analysis contains a (1) preprocessing step (syntactic and ontology tagging), (2) purpose/solution sentence classification and (3) the relationship between two processes is assigned to a control flow (sequential, parallel, causal, etc.).

While those techniques will continue to be essential, new challenges will also emerge as we understand more about the nature of human experience and its role in making an application system. This knowledge grounded in scientific texts may present complex semantic ambiguities. To the best of our knowledge only anaphora resolutions have been studied in procedural texts. However, semantic ambiguities arise more difficulties : *i.e.* word sense ambiguities (polysemy). Word Sense Disambiguation (WSD) [12] is the process to identify the sense or senses of a polysemic terms. Although important, WSD is an ‘intermediate’ task, it is unlikely that anyone other than linguistics would be interested in its results. However, we found that, at least in bioinformatics domain, polysemic words are also common even in very technical domains such as in phylogenetic analyses where specific softwares and programs could be used for different purposes. Generally, programs may contain multiple packages and processes and offer multiple supports. Thus, even specific programs may be used in different contexts. Without a context-based approach it is very difficult to reconstruct a phylogenetic analysis workflow from texts with a simple term-based extraction pipeline. Identifying activities (tasks) and organize them in a control-flow seem to be a complicated task, specially when it can be applied in different positions in a workflow.

We tried to not limit ourselves to highly structured procedural texts but to a domain specific problem to extract only domain-relevant meanings from it. We propose in this paper an ontology-based approach to extract enriched bioinformatics workflows from texts.

Section 2 presents certain choices made in designing our ontology-based extraction process. Two formalisms are presented : workflow representation and ontologies. Additionally, one running example is introduced to explain our representational choices for phylogenetic analyses. Section 3 is the main core of the paper. It describes in details the process of workflow extraction. A text is analyzed with a pipes-an-filter NLP pipeline. Section 4 presents experiments and results of our proposed process by comparing automatically obtained workflows with a gold standard. Section 5 finally discusses results and future work.

## 2. Formalisms for enriched workflow representations

### 2.1. Workflow representation

Traditionally, workflows are "*the automation of a business process, in whole or part, during which documents, information or tasks are passed from one participant to another for action, according to a set of procedural rules*" [1]. In general, workflows consist of a set of *tasks* (activities) combined with a *control-flow* such as sequences, parallel (conjunction) or alternative branches (disjunction), as well as repeated executions (loops). We are interested in this work in workflows as sequences and parallel executions of program tasks. These latter consume *data items i.e.* input sources and output products. The *data-flow* presents interactions (relationships) between them (data items) and tasks. In addition, a task has a set of semantic descriptors (*e.g.* parameters). Data items too. These metadata describe additional information about data, *e.g.* provenace of data.

Different types of workflow representations are proposed in the literature of POCBR [2, 1]. Petri Networks are the most popular design representing the dynamic aspect of processes. However, the representational bias (as in other languages) of such model should be considered while extracting workflows, e.g. multiple instantiations of a task or sub-task. Enriched workflows with semantics describing task parameters and metadata on data items are also to be considered while representing workflows in order to make the workflow more expressive and hence reusable.

In this study, we illustrate our approach in the domain of phylogenetics [13]. Inferred from nucleic acid (DNA or RNA) or protein sequences, phylogenetic relationships represent the evolutionary history of a group of species. During phylogenetic steps, multiple bioinformatics programs (softwares) are used. Here, the tasks represent phylogenetic *programs* and their combination of *parameters* and data items referring to input/output *data* are enriched with *metadata*.

An example of a phylogenetic analysis of hemagglutinin (HA) protein sequences, extracted from the article text of [14], is illustrated in Fig. 1. This workflow represents the phylogenetic solution to study the evolutionary history of HA proteins according to the following steps : (1) multiple sequence alignment, (2) model representation, (3) tree inference and (4) result visualization. Each step is composed of partially ordered program tasks dependently linked with each other. These dependencies represent control and data flows enhanced with semantic annotations. For instance, BionJ, a neighbour joining program, gets Hemagglutinin sequences collected from the Genbank database as input and produces phylogenetic trees.

**Figure 1:**
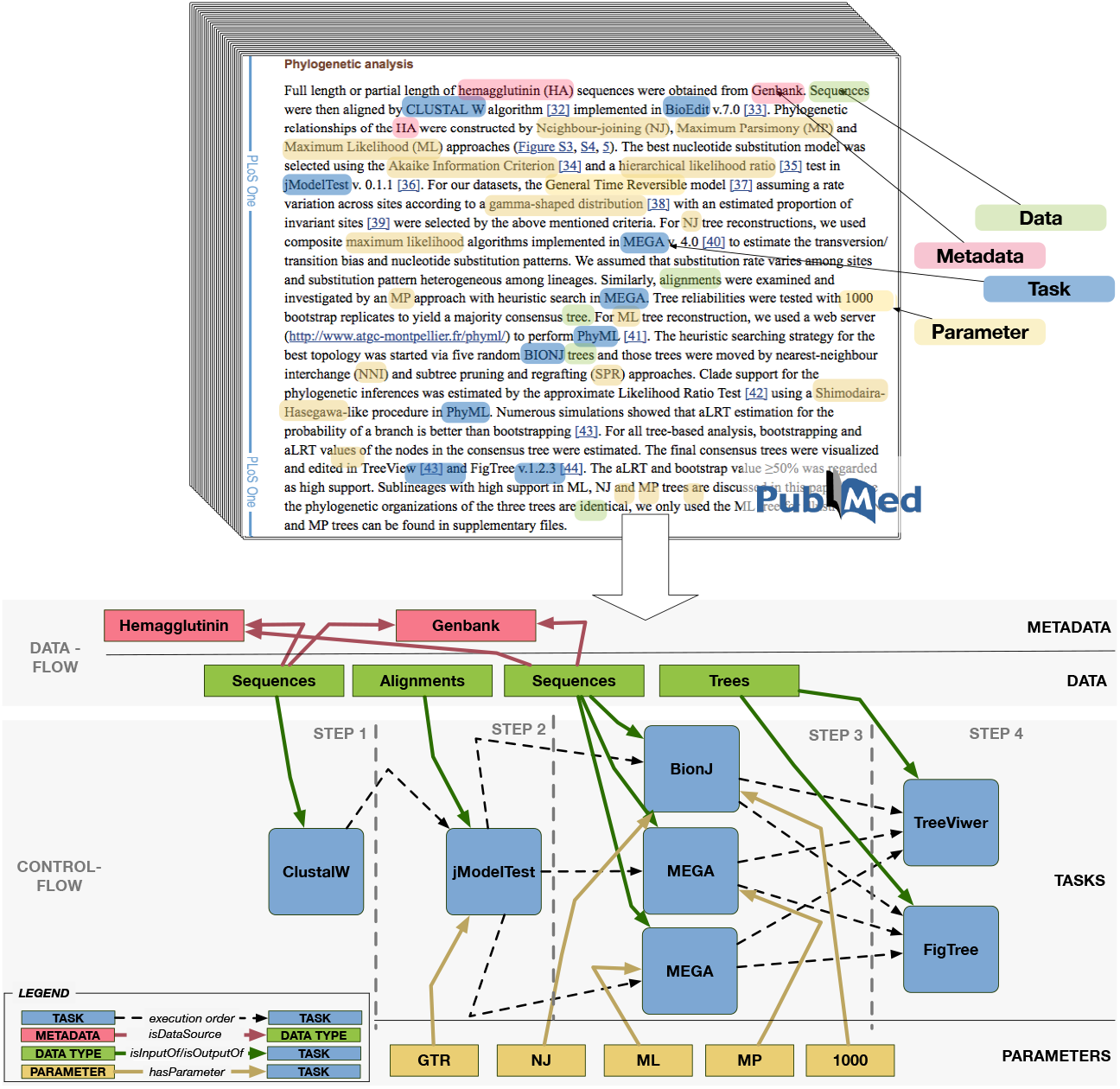
Example excerpts from the article [14].

#### Definition 2.1 (Workflow data language).

*Given a universe of task items and data items* ω^*O*^, *a workflow language* ∆_*W*_ *defines workflow records of partially ordered sequences of inter-related objects o* ∈ *O. A workflow w* ∈ ∆_*W*_ *is a direct acyclic graph* 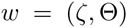 *where* 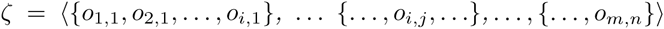 *is the sequence of objects i at each transaction j (step). In turn, 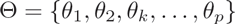 is the set of triples where θ*_*k*_ = (*i*_1_, *l*, *i*_2_) *represents the link of type l between the object at position i_1_ in w.ζ, i.e. w.ζ*[*i*_1_] *and the object at the position i*_2_, *i.e. w.ζ*[*i*_2_].

Intra-transactional relations are not allowed in ∆_*W*_ forbids links within the same transaction : in a valid triple (*i*_1_, *l*, *i*_2_), *i*_1_ and *i*_2_ belong to different transactions whereby the transaction of *i*_1_ is before the one of *i*_2_.

#### Definition 2.2 (Transaction mapping)

*Let* 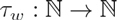 *be the function that maps item positions to transaction positions.* 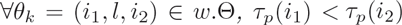.

Fig. 2 shows a sub-workflow representation from the above example in Fig. 1. Phylogenetic programs and parameters and their input/output data and metadata are represented in a partially ordered sequence and a set of relations between them. For example, Hemagglutinin alignments 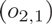 are the input (*θ*_3_) for the neighbour joining program *BionJ* 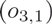 which generates phylogenetic trees (*θ*_4_). The order between phylogenetic programs represents the partial execution order of phylogenetic steps. The parallel execution of programs is represented by transactions in *w.ζ.* Thus, a workflow *w* is represented by a sequence of itemsets (of tasks and data items) and a set of links between them defined in the universe of objects in an ontology.

**Figure 2:**
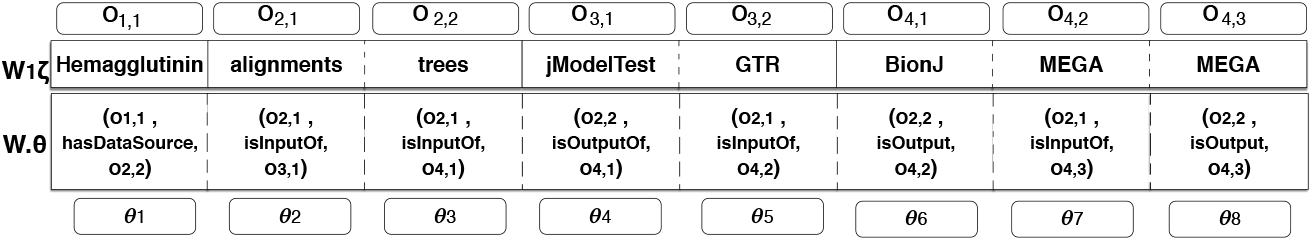
Workflow Sequence Language example as a set of partially ordered concept *w.ζ* and a set of relation triples *w*.Θ. Here, items of the same set are seperated with dotted lines.

### 2.2. Ontology representation

Formally, an ontology is defined as a six-tuple 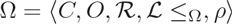 where *O* is the set of all object items, *i.e.* individual tasks and data types, 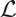 the set of all links between objects, *C* the set of all concepts, and 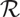 the set of all relations between concepts. Moreover, concepts and relations are organized in taxonomies *w.r.t.* the generality order (*is-a* relationship) of the ontology 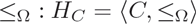 and 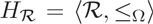, the hierarchical order in the ontology. 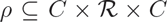 is a ternary relation whose triples 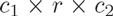 express the connection between a relation and its domain and range concepts (*c*_1_ and *c*_2_, respectively). Objects 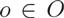 and links 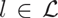 are instances of concepts 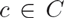 and relations 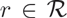, respectively.

#### Definition 2.3 (Universe of objects).

*The universe of objects O^w^ is defined in the domain ontology* Ω. *Formally, a six-tuple 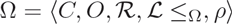 where C is the set of concepts, O is the set of objects, 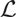 is the set of links and 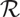 is the set of domain relations. Concepts and relations are organized in taxonomies : 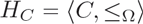 and 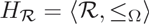, where* ≤_Ω_ *is the generality order (e.g ‘subClassOf’ RDF relationship) in the ontology* Ω. *ρ is a ternary relation 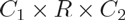 which connects two concepts C_i_ with one relation 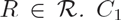 is called the domain concept and C_2_ is the range concept.*

Fig. 3 shows a sample from the domain ontology of phylogenetic anlayses^2^. Grey rectangles represent concepts, hard arcs represent the hierarchical relationships (*i.e.* ‘hasSubClass’ which is the inverse of ‘subClassOf’ in RDF) bet ween concepts and the dotted ones represent the other semantic relations. For example, the first step in a phylogenetic analysis is the ‘data collection step’ which represents program tasks collecting multiple data (’Data Types’ : highlighted concept in green) from different sources ‘Data Source’. Relations between data types and programs are input/output relationships.

**Figure 3:**
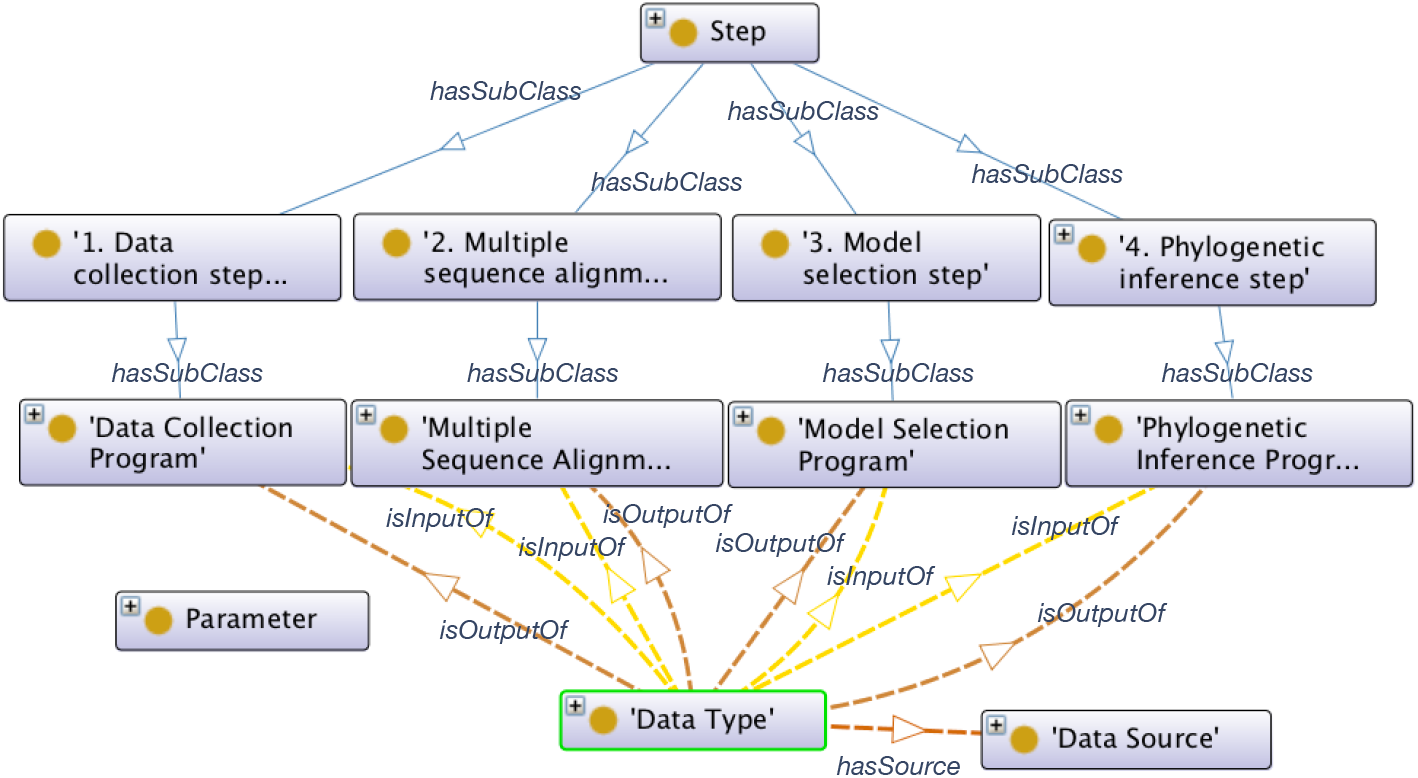
A sample from the phylogenetic domain ontology. Grey rectangles illustrate concepts and coloured arcs represent relationships between them.

## 3. Extraction process

Our framework is based on a pipes and filters architecture [15] using the NLP software GATE (General Architecture for Text Engineering) developed by Cunningham *et al.* [16]. A filter contains an element performing a data transformation on the data stream. The pipes connect the data stream from the output filter to the input one. Filters are independent so one could add or delete them without affecting the execution of others. The following sections describe the different sub-tasks of our workflow extraction solution. Following the standard NLP pipeline, the very first steps are going from the morphological analysis of the text to the identification of syntactic classes of terms and sentences. Section 3.1 quickly goes through those steps. We present some commonly used techniques and tools and we discuss how we adapted them to our issues. Taking the output of those steps, section 3.2 describes our solution to recognize domain terms based on concepts and relations from the ontology. Section 3.3 proposes a way to recognize word sense ambiguities from the ontological concepts. Our WSD technique is supported by a supervised machine learning approach. Section 3.4 describes the way we create a training set to build a WSD and a relational model. Sections 3.5 presents the features and the classifier that we use to learn WSD and relationships in texts. Section 3.6 details the process to reconstruct data and control flows from the extracted and disambiguated terms and finally section 3.7 presents the similarity workflow of extracted workflows based on a gold standard created by an expert.

### 3.1. Morphosyntactic annotation

The initial step segments the text into lexical items called tokens equivalent to words. In general, tokenization operates tokens with white spaces between them as far as we treat english texts. Due to the specificity of the vocabulary of bioinformatics texts, we use a specialized biomedical tokenizer which relies on a dedicated token lattice design pattern and an adapted Viterbi algorithm for bioinformatics tokens [17]. For instance, "*don’t*" is treated as two tokens, while the gene "*4’OMT2*" is really one. The text is also segmented into sentences using punctuation patterns. The extracted tokens and sentences are further tagged with their appropriate grammatical and syntactic classes. At first, a Part-Of-Speech (POS) tagging is executed in order to identify grammatical classes. We use the MedPost POS tagger [18] which is based on a HMM (Hidden Markov Model) model trained over 1,000 XHTML PubMed texts. Their tagger achieved an accuracy of 97.43%. Next, we use a 3-level syntactic parser (*chunker*) to identify active and passive verb phrases in sentences. The depth of the "chunk tree" is strictly limited to 3 (root, phrases and POS tags). Chunkers complexities are equivalent to a finite-state automaton and provides a very simple grammar and sufficient to continue with the extraction process. We present in Fig. 4 an example of a generated chunking tree.

**Figure 4:**
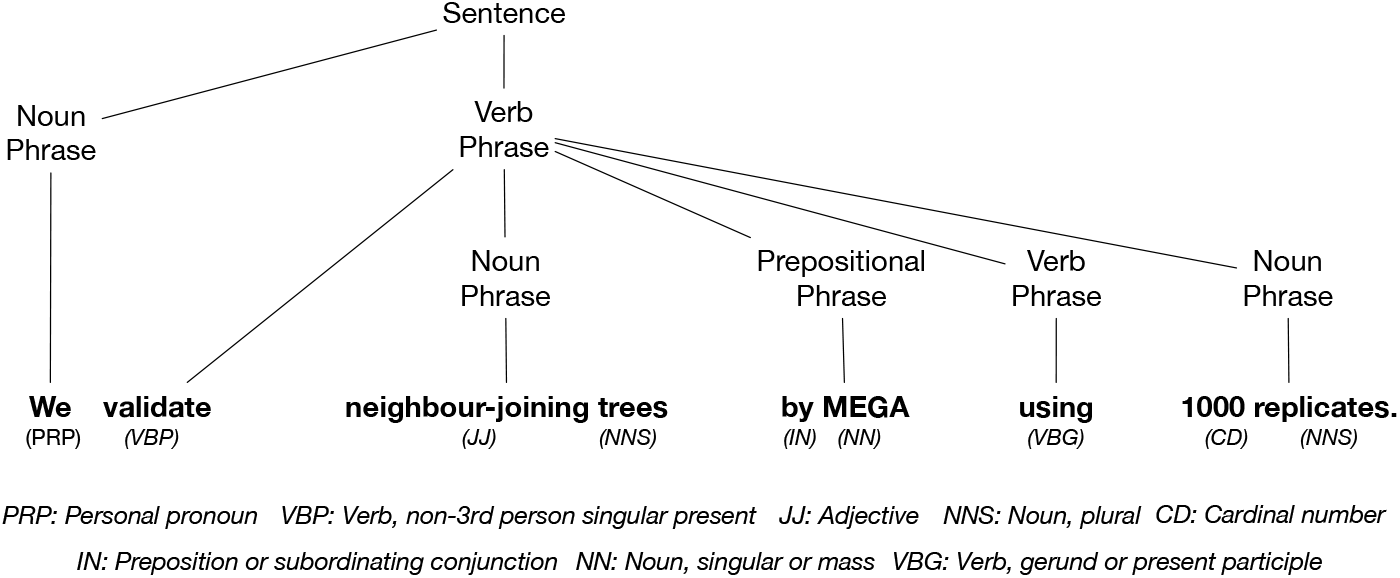
An example of chunking trees.

### 3.2. Semantic annotation

This step is about recognizing workflow items : i.e. tasks, parameters, data, metadata and relations between them. For this purpose, an ontology-based gazetteer (terminology) is used to map domain categories of terms in texts. This step is important to any workflow extraction process. Simple words matchings between ontology instances and text tokens wouldn’t be sufficient to recognize workflow elements in texts. Several syntactic and semantic ambiguities are present in natural language [19]. For this purpose, we use (1) specific JAPE rules to extract and filter workflow concept and relation instances from texts and (2) a word sense disambiguation model to clarify other semantic ambiguities : i.e. multi-class classification (see next sub-sections).

In order to filter concept bad matchings from the ontology, we use specific-domain stop-words, original XHTML markup annotations (from the texts) and regular expression (regex) patterns written in JAPE language. For example, when gene or protein names are nouns, verbs or adjectives, their probability to be exact matchings in a text are never 1. For instance, the gene term ‘*ACG*’ is very similar to DNA codon sequences, *e.g.* in ‘AGC ACT GTA …’. In addition, some bioinformatics software names have similar issues. For example, the word ‘*align*’ in the context of phylogenetics presents a lexical ambiguity too. Commonly, ‘*align*’ is a frequent verb in phylogenetics texts but it’s also used as a program name, that’s why the probability of this term is never 1. However, the probability of ‘align’ to be the name of a software and is written in lowercase is really near to zero. Thus, such pattern is need to be considered. An example of such recognition written in Jape would be :

~~~
 *Rule R*_1_:
({*ClassC.inst* =~ ”*ˆProgramID*_”,
*Token.category == “VB”* || *Token.category == “NN”,*
*Token.orth == “lowercase”*}) : *class*
-->
*inputAS.remove(class)* ;
~~~

Here, *ClassC.inst* is the instance ID in the ontology, *Token.category* is the POS tag of a token and *Token.orth* is its orthography. This rule deletes program terms as verbs and nouns written in lowercase from the set of semantic annotations. Hence, for each (most) specific category^3^ in Ω, we build one rule for the most sensitive POS categories grounding lexical ambiguities : i.e. verbs and nouns.

For the relation recognition step, we adopt the following hypothesis : "*a relation between two concepts (domain and range) may exist in the same sentence evoking the instances of those concepts. The link is defined in the verb of this sentence*". For this purpose, we use ontology triplets *ρ* = (*C*_1_, *R, C*_2_) to match ontological relations *R* with links *L* in texts with *θ* = (*O*_1_, *L, O*_2_). Relation instances (links) are recognized based on all relation triples that exist in Ω compatible with its its elements (domain, range and link type). The interpreter loads all the triples from the ontology and maps tagged concepts as domains and ranges in texts for each link. For example, for the triple (*“BootstrapProgram”, “hasParameter”, “NbBootstraps”*), we search for all its domain and range named entities (instances) from the same sentence in texts. For example, a sentence candidate, in this case, would be : "We validate neighbour-joining trees by MEGA (domain) using (link) 1000 replicates (range)".

If a match is found, a set of patterns is applied for each relation to filter bad matchings. For example, if the phrase voice is active, then the verb evoking the relation domain and range might be between these terms taking into account the order of domain and range terms in the sentence (i.e domain occurring before range). An example of relation pattern recognition is shown bellow (*R*_2_) where *dStart, rStart* and *dEnd* correspond to start and end positions of terms in the sentence.

~~~
*Rule R*_2_ :
(({*Verb Phrase*}) *contains* ({*Domain*} {*Token*}+ {*Range*})) :*p*
-- >
{*for* (*Annotation d* : *domain*)
 *for* (*Annotation r* : *range*)
  *if*(*dStart* < *rStart* && *dEnd* < *rStart* && *p* is *active*)
   *outputAS.add*(*dStart, rEnd,* ”*ClassR*”, *p.verb*)};
~~~

### 3.3. Word sense ambiguity recognition

The previous process of semantic annotation faces known hurdles rooted in the word sense ambiguities : *i.e. polysemy.* The problem is that words often have more than one meaning and sometimes completely different meanings. The meaning of a word in a particular usage can only be determined by examining its context. This is, in general, a trivial task in NLP. For example, in the following two sentences, the software MEGA, a multi-purposes package of programs, is used in different contexts.

— *We validate neighbor-joining trees by MEGA using 1000 replicates.*
— *Tree data was visualized using MEGA.*

In the first sentence, MEGA is used to validate the phylogenetic hypothesis which corresponds to the ‘*HypothesisValidationProgram’* concept. However, in the second sentence, the program MEGA is used in the visualization step which corresponds to the ‘*TreeVisualizationProgram*’ concept. Multi-purpose packages programs such as MEGA, PHYLIP and BEAST refer to multiple concepts in the domain ontology. A new concept ‘*GeneralPurposePackages*’ is added to the ontology Ω. This is done automatically while interpreting the hierarchy of concepts *H*_*C*_ to search for individuals (instances) belonging to multiple task concepts. For each concept *c ∈ C* belonging to a multiple inheritance in the hierarchy *H*_*C*_, we add the RDF triples : 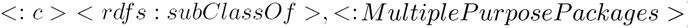 *MultiplePurposePackages* >) in Ω. Moreover, we add the equivalence relationship between *c* and the ambiguous concept ‘*MultiplePurposePackages*’, i.e : 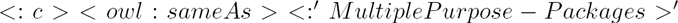 (see example in *Fig.* 5). Hence, after discovering polysemic concepts in the ontology, we tag their ambiguous instances in texts.

**Figure 5:**
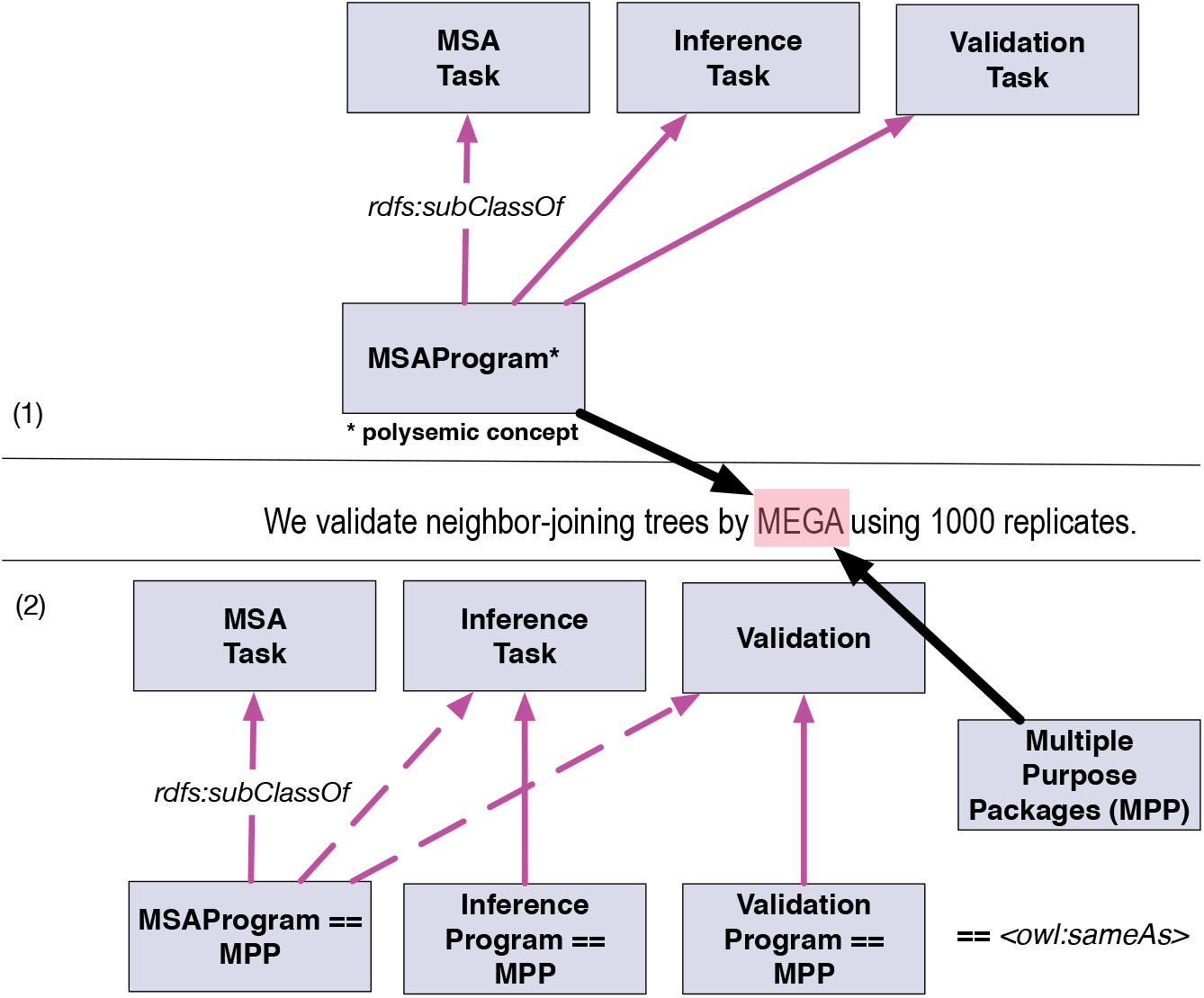
Ontology transformation. An example of multiple inheritance grounding polysemic concepts in Ω : (1) the ontology before (having multiple inheritance), (2) the ontology after (keeping only the most right relation (dotted arrows) and adding a new concept for each ambiguous task).

### 3.4. Word sense disambiguation

We propose here a supervised machine learning approach to WSD. The problem of WSD is defined as follows : "*given a set of pre-tagged polysemic words, WSD is to classify each word with its appropriate task concept (sense) with high accuracy and efficiency*". Pre-tagged words are usually created manually, but here we propose to construct the WSD ground truth in a semi automatic way using the domain ontology and the syntactic tags in sentences.

~~~
*Rule R*_3_ :
*Priority* : 90
(({*VerbPhrase}) contains* ({*GeneralPurposePackages*})) : *context*
-- >
{*if* (*context is active* &&
 *context.verb.root in* {”*valid*”, ”*test*”,…} &&
 *context.object.root in* {”*bootstrap“,”replicate*”, …})
 *GeneralPurposePackages.put*(*”resolution”*, *”ValidationProgram”*);} ;
~~~

Rule *R*_3_ shows an example of JAPE rules to generate the WSD learning set. Here, both of the context concepts (see section 3.2) and the syntactic tree (see section 3.1) are used to clarify the concept c into the ‘*ValidationProgram*’ concept. The set *context.verb. root* are root verbs extracted automatically from the Wordnet synonyms [20] of the parent concept *c*′ in *H*_*C*_ (i.e from the level immediately above : 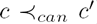). The set *context.object.root* are also Wordnet synonyms of *Parameters* terms. These latter are used (if they exist) as hints to classify the polysemic program to its right class. The pattern *R*_3_ verifies if the verb in the active voice phrase is about testing or validating something. In addition, if a validation parameter is found in the object of the sentence, then the ‘*GeneralPurposePackages*’ concept is disambiguated to the concept ‘*ValidationProgram*’. Each WSD rule has a priority probability to fire them by order. For each polysemic concept created in the previous step, we generate one WSD rule. An additional rule is added by the expert to represent the default concept if no heuristic is valid (default rule).

### 3.5. Learning module

In order to create a model for automatic WSD, a set of features is generated representing the dataset of already tagged words in texts. For example, in order to classify the term ‘MEGA’ (*ClassC*) in the example of Fig. 6, we generate the set of features (*featuresC*) reflecting its context. The context covers a window made of : the *Win*_*i*_ concepts (*Token.class*) and POS tags (*Token.category*) preceding and the *Win*_*i*_ ones following the concerned word in the sentence. The window of concepts and POS tags (*Win*_*i*_) is fixed in the learning step. In this example, *win*_*i*_ = 3, hence, *Token.class = {DataType, MaximumParsimonyApproach}* and *Token.category = {Determiner, Noun, Preposition}.* In addition, in order to classify a relation *(classR=has_used_by_program)* between the ambiguous word ‘MEGA’ (relation’s domain) and the data type ‘alignments’ (its range), we define its context as the set of tokens between the domain and the range. Now, features (*featuresR*) represent context tokens’ POS tags (*Tokens-Between.category*), their distance (in number of tokens between domain and range), direction (domain-to-range) and verbs between the concerned terms. As a feature selection approach, we use a sequential forward selection [21] while adding *f* features at a time. The selected best set of features are then used in the construction of a model for automated annotation of ambiguous words in texts and their relations.

**Figure 6:**
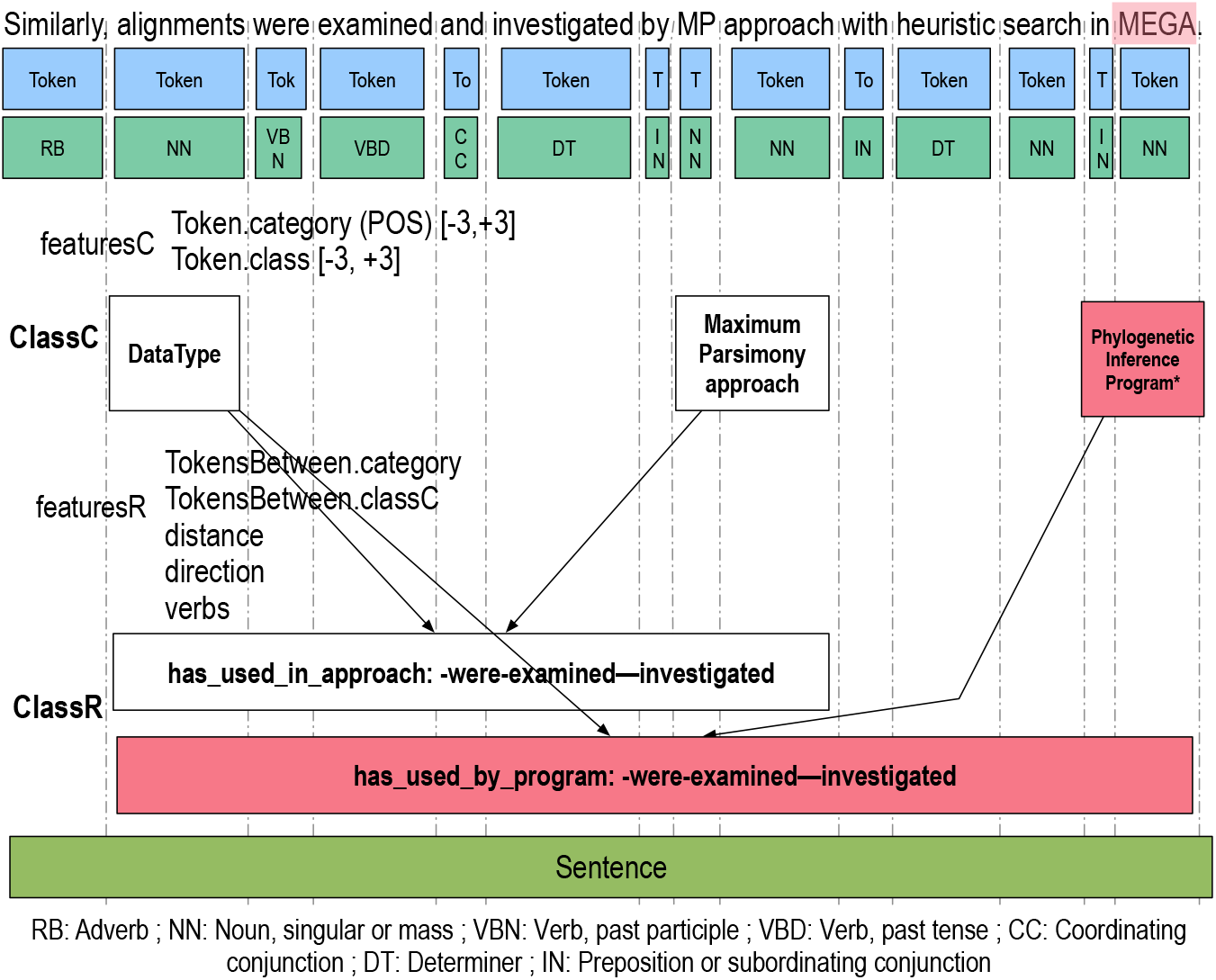
An example of learning features for WSD concepts and relation extraction from texts.

We use the PAUM model (Perceptron Algorithm with Uneven Margins) [22] to learn WSD and relation classifications based on the extracted features. PAUM is a hybrid SVM (Support Vector Machine) and Perceptron (neural network) classifier that was designed especially for imbalanced data and has successfully been applied to various semantic annotation problems [23, 24]. This algorithm differs from other classifiers as it treats positive and negative examples differently by introducing two margin parameters *τ*_+_ and *τ*_−_ into the updating rules for the positive and negative examples, respectively. Here, *τ*_+_ and *τ*_−_ simulate context examples (see *Fig.* 6), hence, we could increase/decrease the distance between an ambiguous concept and its surrounded ones. This reflects the intuition that the nearer a neighbouring concept is, the more important it is for classifying the given token. It is about the same intuition for the relation classification. We modelize the distance between domain and range concepts of one relation with *τ*_−_ and *τ*_+_ parameters representing the positive and the negative token annotations between relation components. PAUM learns how to update its learning rate and margin parameters to maximize this distance [22]. After tuning (in number of features, margin parameters, *etc.*) and training our models, we applied them to the test set (test corpus) which constitutes 1/3 of the available corpus (We note the other 2/3 are used for training). Extracted terms and links are then evaluated using different measures like precision, recall and F1measure [25].

### 3.6. Workflow reconstruction

From recognized links between concept instances in texts, we build our concrete workflow knowledge database where items are the named entities ordered by the top-level pattern in the domain ontology. Concept and relation instances serve to identify respectively the data-flow and the control-flow. For example, from the tuples : *ρ*_1_ = (Alignments, "isOutputOf", ClustalW), *ρ*_2_ = (Alignments, "isInputOf", MEGA), we construct the control-flow : *w.ζ* = 〈{Alignments}, {ClustalW}, {MEGA} 〉 and the data-flow : *w.θ* = (*ρ*_1_, *ρ*_2_, *ρ*_3_) where *ρ*_3_ = (Alignments, "genInputTo", ClustalW). The partial order in the sequence *w.ζ* is carried out with the ‘generate input to’ (*genInputTo*) relation which is the result of the transitive relation between an input and output data between two tasks from the ontology Ω.

*Fig.* 7 shows an example of workflow reconstruction from a domain ontology. The hard black arcs represent the possible concept matchings between the ontological workflow pattern (grey rectangles) and the workflow sequence *w*_1_. *ζ* (blue rectangles). The first set in *w*_1_.ζ : {Sequences, Alignments, Tree} represents the data types. While relations *ρ* between datatypes and programs ‘ClustalW’ and ‘MEGA’ represent the data-flow, the sequence of itemsets in *w*_1_.*ζ* represent the control-flow.

**Figure 7:**
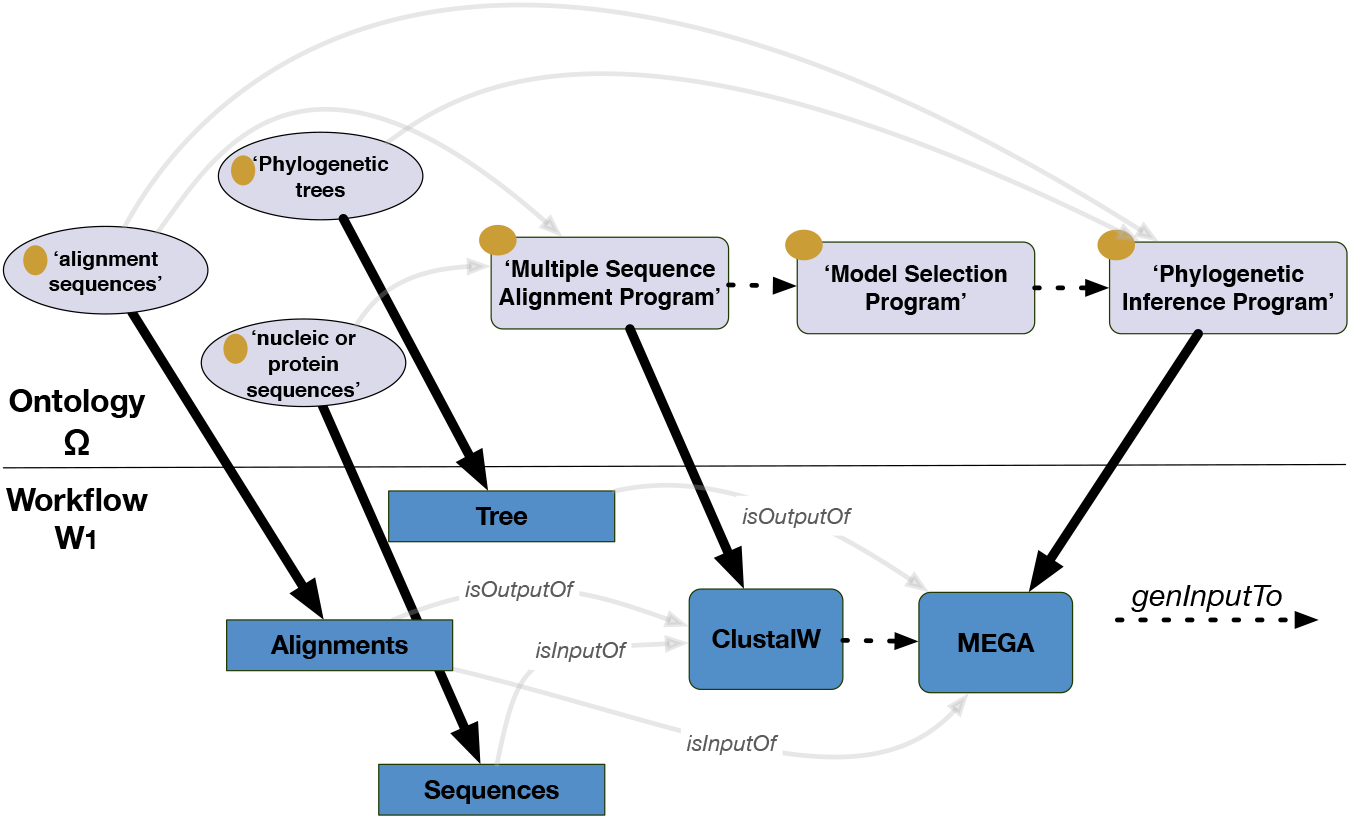
An example of a workflow reconstruction from a domain-ontology.

### 3.7. Workflow similarity measure

Here, we calculate the similarity measure *s*_*w*_(*a,p*) ∈ {0…1} which calculates the similarity between a gold standard workflow *a* and a predicted one *p.* Each workflow *a, p* is represented by the couple (*ζ*, Θ) in ∆_*W*_. The similarity between two workflows *a* and *p* is given by the following formulas :

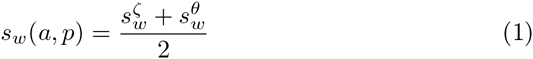

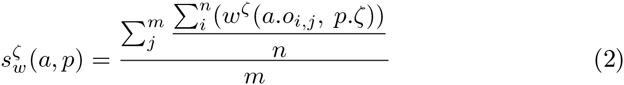

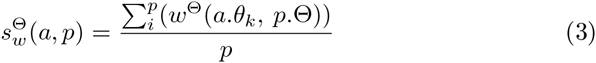

Here *m*, *n* and *p* are respectively the total number of steps, items per step and relations in a workflow. Each object *o*_*i,j*_ is weighted by the accuracy measure 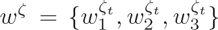 and each property *θ*_*k*_ is weighted by 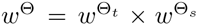 where the weight *w*^Θ_*t*_^ ∈ {0…1} is the degree of an object term accuracy and *w*^Θ_*s*_^ ∈ {0..1} is the degree of a link term accuracy. The semantic similarity of a predicted workflow is then the mean of its component accuracies in the gold standard.

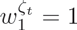 is the weight given if the object *a.o*_*i,j*_ is found in *p.σ* and is correct. 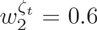 is the weight given if the object *a.o*_*i,j*_ is found in *p.σ* and is partially correct. Finally, 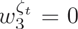 if the object *a.o*_*i,j*_ is not found in *p.σ.* In the relation side, we tuned the weights as follows : 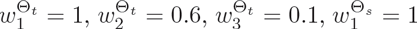, 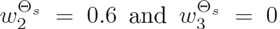. Hence, *w*^Θ_*t*_^ are the weights given if the term of a link (*e.g. ‘isDataSource’*) is correct, partially correct or not. Same for 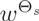, weights represent the correctness degrees of relations structure : domain and range terms. Weights values are inspired from the work of [26] where lenient precisions and recalls are calculated given a partial overlap measure of terms.

## 4. Experiments and results

### 4.1. Dataset

For this study, we downloaded scientific articles from the digital repository PubMed Central PMC database (published from January until April 2015) in XHTML format. 320 texts have been extracted. At first we downloaded more than 1000 recent articles, journals and conference papers having at least one section : ‘phylogenetic analysis’ or ‘phylogenetic analyses’. However, these sections may have not sufficient information to reconstruct workflows from texts. Thus, we filtered the downloaded articles based on the smallest ontological pattern that any phylogenetic analysis should represent. A phylogenetic study might use at least one datatype and a phylogenetic inference program (see Fig. 3).

In addition, we constructed at first a relational database gazetteer (in MySQL) from different well-known databases in the phylogenetic literature such as Gene ontology, NCBI, UniprotKB and from Felsenstein’s web site^4^ [27]. The terminology is then reorganized in a domain ontology in order to recognize terms and relations in texts. We used the Quest-Ontop plateform [28] to map MySQL classes to SPARQL concepts in a domain ontology. The generated ontology presents 17 different object properties (relations), 44,581 concepts (with 7 root concepts and 111 unique concepts) and more than 1,300,000 unique instances without counting the synonyms. Root concepts represent the seven top tasks (steps) of a phylogenetic analysis. Our ontology is free to access under the Bioportal repository : http://bioportal.bioontology.org/ontologies/PHAGE. The Gate implementation of our workflows extractor is available in : http://labo.bioinfo.uqam.ca/tgrowler.

### 4.2. Results

Following experiments are obtained from the 100 extraction results by the human expert (gold standard).

In the first experiment, we evaluate the PAUM WSD model. We use precision, recall and F1measure metrics. A 10-fold cross validation was run over the PAUM classifier to choose the best parameters : 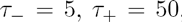, where *τ*_−_ and *τ*_+_ are respectively the negative and positive margins. We have picked the best values for *τ*_−_ from {−10, −5, −1,0, 1,5, 10} and for *τ*_+_ from {−100, −50, −10,0, 10, 50, 100} based on the validation test. We chose the whole set of proposed features, since all POS and concept context tags give the best F1Measure. The best context window is *win*_*i*_ = 5 based on our experiments.

In the second experiment, we evaluate the relation extraction model. Same as in the previous experiment, we ran a 10-fold cross validation over the PAUM Relation Extraction (PAUM RE) classifier. We chose the best features and best parameters as follows : *τ*_−_ = 1, *τ*_+_ = 10. Selected features are : tokens POS, concepts, distance (in terms of number of tokens), direction and the list of verbs between domain *d* and range *r* links. Due to the space constraints, we only show here the precision, recall and F1Measure for the best configurations.

Table 1 shows results of WSD and relation extraction learning. For PAUM WSD, the recall is the ratio of total amount of ambiguous words correctly disambiguated over the total amount of ambigious words in the gold standard, and the precision is the ratio of the total amount of correctly disambiguated words over the total of generated ambigious words. Our PAUM WSD model is highly precise (99.21%) but predicted words represent 66.66% of the total number of ambiguous words in the gold standard. This is understandable since the context of an ambiguous word could be out of it’s sentence context. Hence this is one of our context-based limitation hypothesis. PAUM RE model presents different results in terms of precision and recall from the PAUM WSD but it shows similar F1Measures. PAUM RE covers 84.22% from the actual relations in the gold standard, however, 72.63% of the generated relations are correctly predicted.

**Table 1:** PAUM Word Sense Disambiguation and relation extraction evaluation results.

|  | Precision (%) | Recall (%) | F1Measure (%) |
| --- | --- | --- | --- |
| PAUM WSD | 99.21 | 66.66 | 79.74 |
| PAUM RE | 72.63 | 84.22 | 77.24 |

Fig. 8 shows the similarity distribution results obtained through the evaluation of 100 gold standard workflows. Generated workflows are 82.89% similar with the actual workflows extracted by the expert (first figure on top in Fig.8). While, the best mean workflow similarity is 100%, the worst workflow similarity is 54.68% which means that at the worst cases our workflow extraction framework will probably extracts the half of the real workflow components.

**Figure 8:**
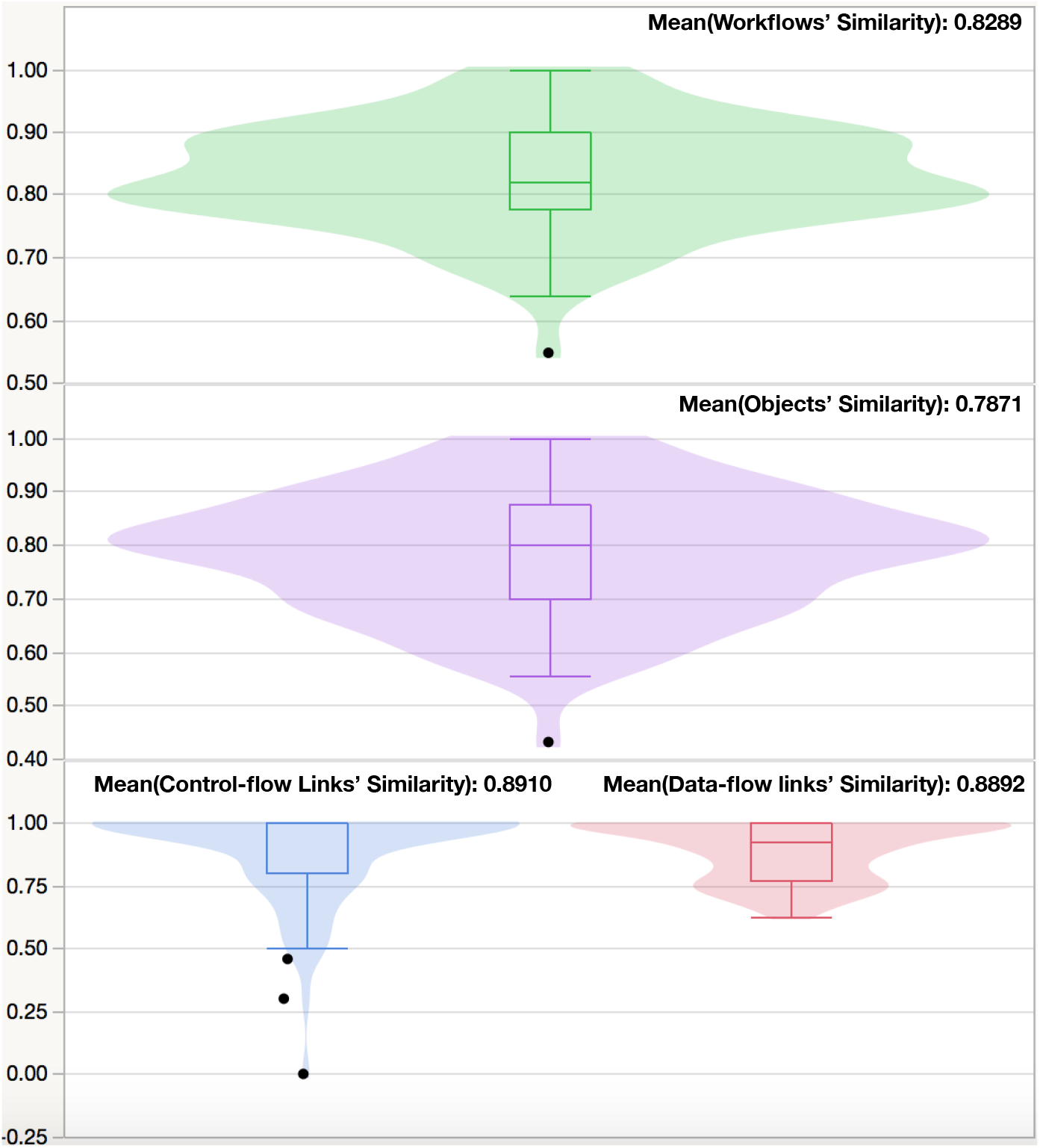
Workflow similarity distributions.

The mean of workflow object similarities is 78.71% (second figure in Fig.8). The best workflow task similarity is found in phylogenetic tree inference programs (100%). Generally, this task is explicitly defined in phylogenetic analysis texts. We found that phylogenetic methods are frequently co-occurred in the context of the phylogenetic inference programs which explains the mean similarities of this task. However, the worst task similarity, ‘Hypothesis Validation’, is about 60%. This is due to the context of this task which is ambiguous to be be resolved in one sentence. 30% of the hypothesis validation programs are ambiguous programs (multiple purpose packages). Concepts and POS categories surrounding these programs are not very conclusive to disambiguate this category of tasks.

The relation side of workflows rep resents the data-flow and control-flow (third figure in Fig.8). The mean similarity in the control flow is 89.1% and 88.92% in the data flow. While the worst similarities in the data-flow relations is 74.43% which represents the relation between data types and multiple sequence alignment programs, the best data-flow similarity is the similarity of the relation between data types and hypothesis validation programs (100%). Verbs like "validate" and "test" are commonly used between data types and these programs which explains the high similarity in this concept. The worst similarities in the control-flow are found in the links between phylogenetic inference programs and their methods with a mean of 79.84%. However, links between programs’s parameters like the number of bootstraps and the type of evolutionary models are, generally, explicitly described in the sentence which explains its high similarity in the control-flow with a mean of 100%.

### 5. Conclusion

This paper addresses the problem of the extraction of enriched workflows using a domain-ontology to resolve word sense ambiguities in texts. The framework is supported by an application in bioinformatics domain. Our graph based workflow representation extends the classic POCBR workflow structures with more semantic relations that serve for both : the extraction and the resolution of polysemic tasks in texts. It’s promising that our method reaches high similarities with an ontology-based word sense disambiguation approach for tasks, however, other concept ambiguities need to be improved, mainly data type metadata such as : genes, proteins and species terms.

Future work will extend the existing control-flow representation with more sophisticated relations representing disjunction and loop controls in workflows. Outside the context of the case acquisitions, we intend to include text adaptation where, in the case of extraction deficiencies (missing tasks), fragments from the workflows are automatically added based on more general contexts in texts instead of sentences : *e.g.* paragraphs.

## Footnotes

1. Since 1880.

2. http://bioportal.bioontology.org/ontologies/PHAGE

3. A most specific concept/relation is the leaf category (which has no child category) in the hierarchies *H*_*C*_ and *H*_*R*_.

4. http://evolution.genetics.washington.edu

